# Kinetic evidence for multiple aggregation pathways in antibody light chain variable domains

**DOI:** 10.1101/2023.08.28.555139

**Authors:** Sherry Wong, Madeline E. West, Gareth J. Morgan

## Abstract

Aggregation of antibody light chain proteins is associated with the progressive disease light chain amyloidosis. Patient-derived amyloid fibrils are formed from light chain variable domain residues in non-native conformations, highlighting a requirement that light chains unfold from their native structures in order to aggregate. However, mechanistic studies of amyloid formation have primarily focused on the self-assembly of natively unstructured peptides, and the role of native state unfolding is less well understood. Using a well-studied light chain variable domain protein known as WIL, which readily aggregates *in vitro* under conditions where the native state predominates, we asked how the protein concentration and addition of pre-formed fibril “seeds” alter the kinetics of aggregation. Monitoring aggregation with thioflavin T fluorescence revealed a distinctly non-linear dependence on concentration, with a maximum aggregation rate observed at 8 µM protein. This behavior is consistent with formation of alternate aggregate structures in the early phases of amyloid formation. Addition of N- or C-terminal peptide tags, which did not greatly affect the folding or stability of the protein, altered the concentration dependence of aggregation. Aggregation rates increased in the presence of pre-formed seeds, but this effect did not eliminate the delay before aggregation and became saturated when the proportion of seeds added was greater than 1 in 1600. The complexity of aggregation observed *in vitro* highlights how multiple species may contribute to amyloid pathology in patients.

## INTRODUCTION

Aggregation of normally soluble proteins as amyloid fibrils and other non-native species leads to progressive tissue damage in systemic amyloid diseases (Muchtar et al., 2021; Buxbaum et al., 2022). In amyloid light chain (AL) amyloidosis, the amyloid fibrils are derived from monoclonal antibody light chain proteins, which are secreted from an expanded population of clonal plasma B cells (Merlini et al., 2018; Morgan et al., 2021). Each individual with AL amyloidosis has a light chain with a unique sequence that forms their amyloid fibrils, and this sequence diversity may contribute to the clinical heterogeneity of disease (Enqvist et al., 2007; Bodi et al., 2009; Kourelis et al., 2017). Of the two types of human light chains, kappa and lambda, AL amyloidosis is more frequently associated with lambda light chains, while kappa light chains are more frequent in the healthy immune repertoire. How and why certain light chain sequences can form amyloid in patients is not understood, and mechanistic details on the process of amyloid formation could contribute to efforts to treat the disease.

Antibodies are secreted as globular folded proteins, usually hetero-oligomers of heavy chains and light chains, but excess light chains can be secreted without a heavy chain partner (Feige et al., 2010). These “free light chains” can form homodimers that may be stabilized by an inter-chain disulfide bond. Light chains have two structural immunoglobulin domains, an antigen-binding variable domain and a constant domain. Each domain has a ß-sandwich fold, stabilized by a single intramolecular disulfide bond. AL amyloid fibrils are most commonly formed from fragments of light chains, including the variable domain (Connors et al., 2007; Lavatelli et al., 2008; Enqvist et al., 2009). High resolution structures of amyloid fibrils show that their cross-ß core is formed from variable domain residues in non-native conformations, with the disulfide bond intact but the orientation of the peptide chains around the disulfide reversed, relative to that in the native state (Radamaker et al., 2019, 2021b, 2021a; Swuec et al., 2019). Unfolding of the native light chains is therefore required for amyloid formation. Consistent with this observation, aggregation rates of folded proteins, including LCs, are correlated with the stability of their globular domains (Hurle et al., 1994; Wall et al., 1999; Souillac et al., 2002a; Rennella et al., 2019b).

Amyloid formation has been successfully modeled as a nucleated polymerization reaction, where slow formation of an initial nucleus is followed by rapid extension by addition of molecules to the growing fibril (Ferrone, 1999; Powers and Powers, 2008; Meisl et al., 2016). Measuring aggregation kinetics as a function of protein concentration, temperature or other factors has revealed details of how protein sequence influences aggregation. However, this framework was designed to describe aggregation of unstructured peptides where amyloid-competent conformations are accessible to all soluble proteins at equilibrium. It is not clear how the two potential fates of an unfolded protein – folding to the native structure or assembly to an aggregate structure – are resolved under conditions where both conformations are accessible (Morgan, 2022).

To gain insights into the mechanisms of light chain aggregation and potential roles of nucleation, we here investigate the aggregation kinetics of a lambda light chain variable domain known as WIL, which was originally identified in a patient with AL amyloidosis (Wall et al., 1999). WIL is derived from the germline antibody precursor gene *IGLV6-57*, which is the most common precursor gene observed in AL amyloidosis clones (Bodi et al., 2009). The isolated variable domain of WIL readily forms amyloid fibrils *in vitro* at neutral pH and 37°C. We used binding of the fluorogenic dye thioflavin T (ThT) to measure the early stages of aggregation and asked how the presence of pre-formed seeds and the addition of peptide tags to the protein N- or C-termini alters the rate at which ThT-binding aggregates are formed.

## RESULTS

### WIL-V aggregation has a non-linear dependence on concentration

We measured the aggregation kinetics of a well-studied light chain variable domain known as WIL (Wall et al., 1999). The original paper studied the isolated WIL variable domain, which we call WIL-V to distinguish it from a full-length LC construct, WIL-FL, where an *IGLC3* constant domain was fused to the previously-published variable domain sequence (Morgan and Kelly, 2016). Recombinant WIL-V was expressed in the *E. coli* periplasm without a fusion tag. Although WIL-V can form homodimers, it does so with a dissociation constant of around 5 mM (Rennella et al., 2019a), so at the concentrations used here (1-32 µM) it is predominantly monomeric. As previously observed, incubation of WIL-V at 37 °C in phosphate-buffered saline (PBS, pH 7.4) with agitation in microwell plates leads to formation of aggregates that bind to ThT (Wall et al., 1999; Rennella et al., 2019a). When the aggregation reactions are started, ThT fluorescence remains low for a “lag phase” of several hours, after which it increases rapidly in a “growth phase” to reach a plateau (Figure 1a). Further incubation leads to a decrease in ThT fluorescence (Supplemental Figure 1) that we attribute to either conformational changes in the aggregate or bleaching of the ThT (Morgan and Kelly, 2016). Under these conditions, the full-length protein WIL-FL remains soluble, as we have previously reported (Supplemental Figure 1) (Morgan and Kelly, 2016).

**Figure 1:**
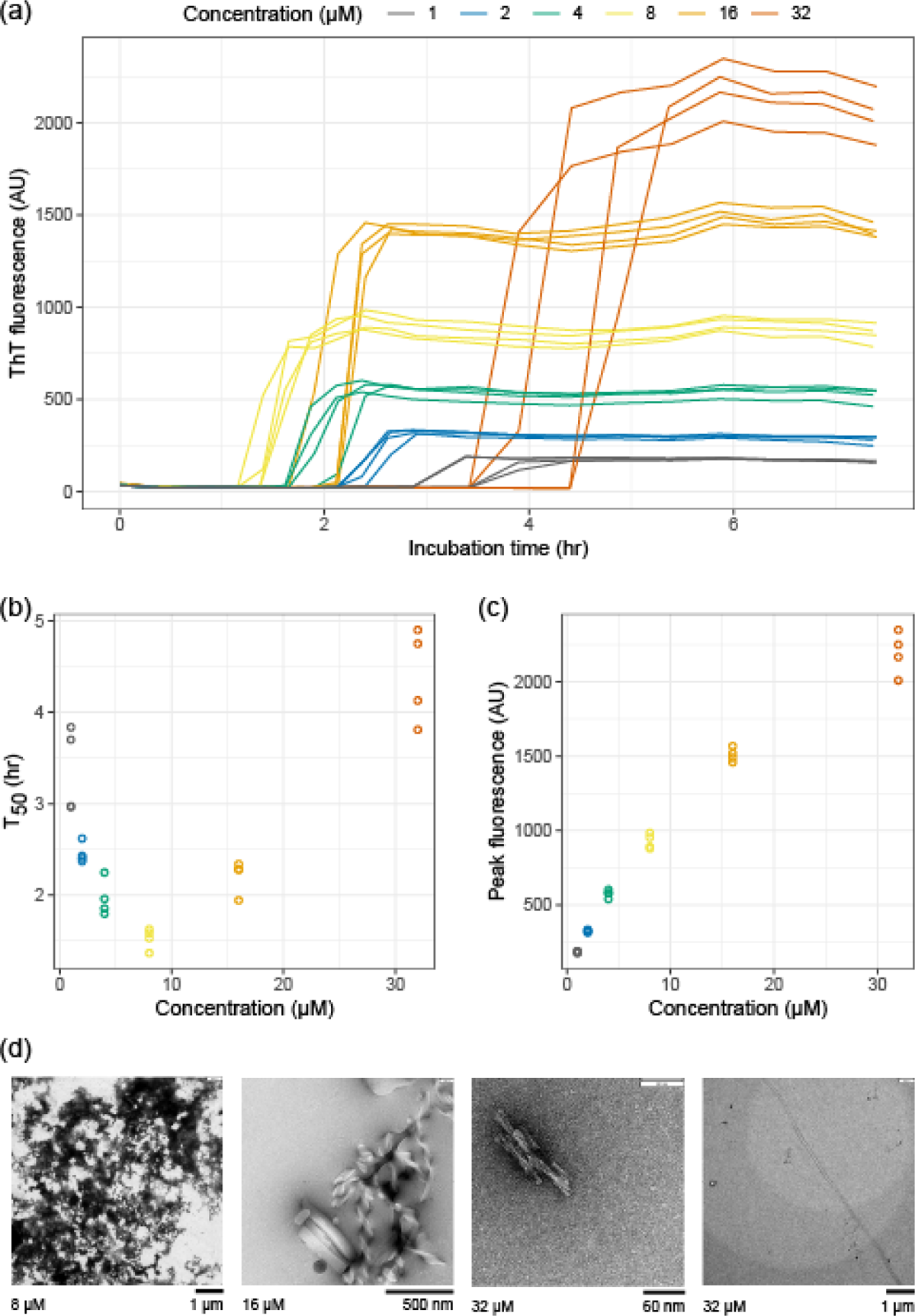
Aggregation kinetics of WIL-V have a nonlinear dependence on concentration. WIL-V solutions were incubated in microplates in PBS, pH 7.4, containing 1 µM ThT at 37 °C and shaken at 500 rpm. Data shown from two independent microwell plates, each with two wells per condition. (a) ThT fluorescence as a function of time. Colors indicate initial WIL-V concentration. (b) Calculated aggregation midpoint (T_50_) as a function of concentration from the data shown in (a). (c) Maximum fluorescence intensity from the data shown in (a). (d) Electron micrographs showing examples of aggregate morphology observed after 72 h of incubation. Initial WIL-V concentrations and appropriate scale bars are shown below each image. Multiple aggregate species were observed in each reaction.

Monitoring ThT fluorescence kinetics as a function of WIL-V concentration reveals a chevron- or V-shaped dependence on concentration (Figure 1b). The length of the lag phase decreases between 1 µM and 8 µM WIL-V, then increases at higher concentrations. This observation is consistent with the rate of ThT-positive aggregate formation being limited by accumulation of non-ThT-binding species. However, the peak ThT fluorescence was proportional to the initial concentration of WIL-V, consistent with all protein converting to amyloid-like material during the rapid growth phase of the reaction (Figure 1c). Negative-stain electron micrographs of aggregates (Figure 1d) show a mix of amorphous deposits, ribbon-like structures and short fibrillary material, but few of the long amyloid fibrils that we have previously observed for WIL-V and other variable domains after longer incubation times (Morgan and Kelly, 2016; Rennella et al., 2019b).

To gain further insight into the molecular changes that underlie these kinetics, we attempted to fit the traces individually and globally to models of nucleated aggregation. We have not identified a mechanistic model that can robustly fit these data, likely because these models were developed to describe systems where unstructured peptides aggregate in quiescent reactions. For example, Supplemental Figure 2 shows the output from an unsuccessful attempt to use the AmyloFit system (Meisl et al., 2016) to fit the WIL-V data to a nucleation-elongation model. We therefore used a non-mechanistic sigmoidal model fit to extract T_50_ values, the time at which the reaction reaches its midpoint fluorescence, for each individual reaction.

### Modulation of WIL-V assembly without altering stability

The rapid increase in ThT fluorescence characteristic of amyloid formation is attributed to elongation of fibrils by soluble precursors once an initial nucleus has formed, and the lag phase is interpreted as the time required for sufficient nuclei to form. Extension of amyloid fibrils involves precise structural templating by the fibril, and requires the soluble protein to adopt a conformation consistent with addition to the fibril. Therefore, we hypothesized that structural changes in the precursor protein or fibrils might alter the ability of fibrils to form or extend nuclei. However, previous investigations into light chains show that the stability of the native state is a major determinant of aggregation rate, and that fibrils appear to tolerate mutations that might be expected to disrupt their structure (Rennella et al., 2019b). To investigate the roles of nucleation and extension in WIL-V aggregation without greatly altering the folded state stability, we asked how addition of N- and C-terminal peptide extensions alter the aggregation kinetics. Such extensions should not greatly alter the equilibrium between the folded and unfolded states of the protein. However, they may enhance or disrupt interactions within the aggregates, or between the unstructured species that must self-assemble to form amyloid. We added N- or C-terminal histidine tags since these were readily accessible by cloning and could be used to aid purification. We refer to these proteins as WIL-V-Nhis and WIL-V-Chis, respectively. For many proteins, the presence of a his-tag does not greatly alter folded domain structure or stability (Gorski et al., 2001). Histidine residues are predominantly uncharged at pH 7.4 and able to form cross-ß structures. However, an N-terminal extension is incompatible with the only known structures of amyloid fibrils formed by a light chain known as AL55, which is derived from the same *IGLV6-57* precursor gene as WIL-V (Swuec et al., 2019; Puri et al., 2023). In contrast, AL amyloid fibrils often comprise residues from the light chain constant (Olsen et al., 1998). Both N- and C-terminal extensions are both compatible with other known AL fibril structures. We verified that our three WIL-V constructs form predominantly folded structures at 37 °C in PBS (Figure 2a) and have similar urea unfolding midpoints (Figure 2b), as assessed by intrinsic tryptophan fluorescence. WIL-V has a single tryptophan residue that packs close to the disulfide bond in the native state, resulting in low fluorescence intensity. In the unfolded state, the tryptophan residue has a higher fluorescence with an emission maximum of 355 nm. Because these proteins have similar folded structures and stabilities, we expected that differences in their aggregation behavior would be attributable to the effects of their his-tags on aggregate self-assembly.

**Figure 2:**
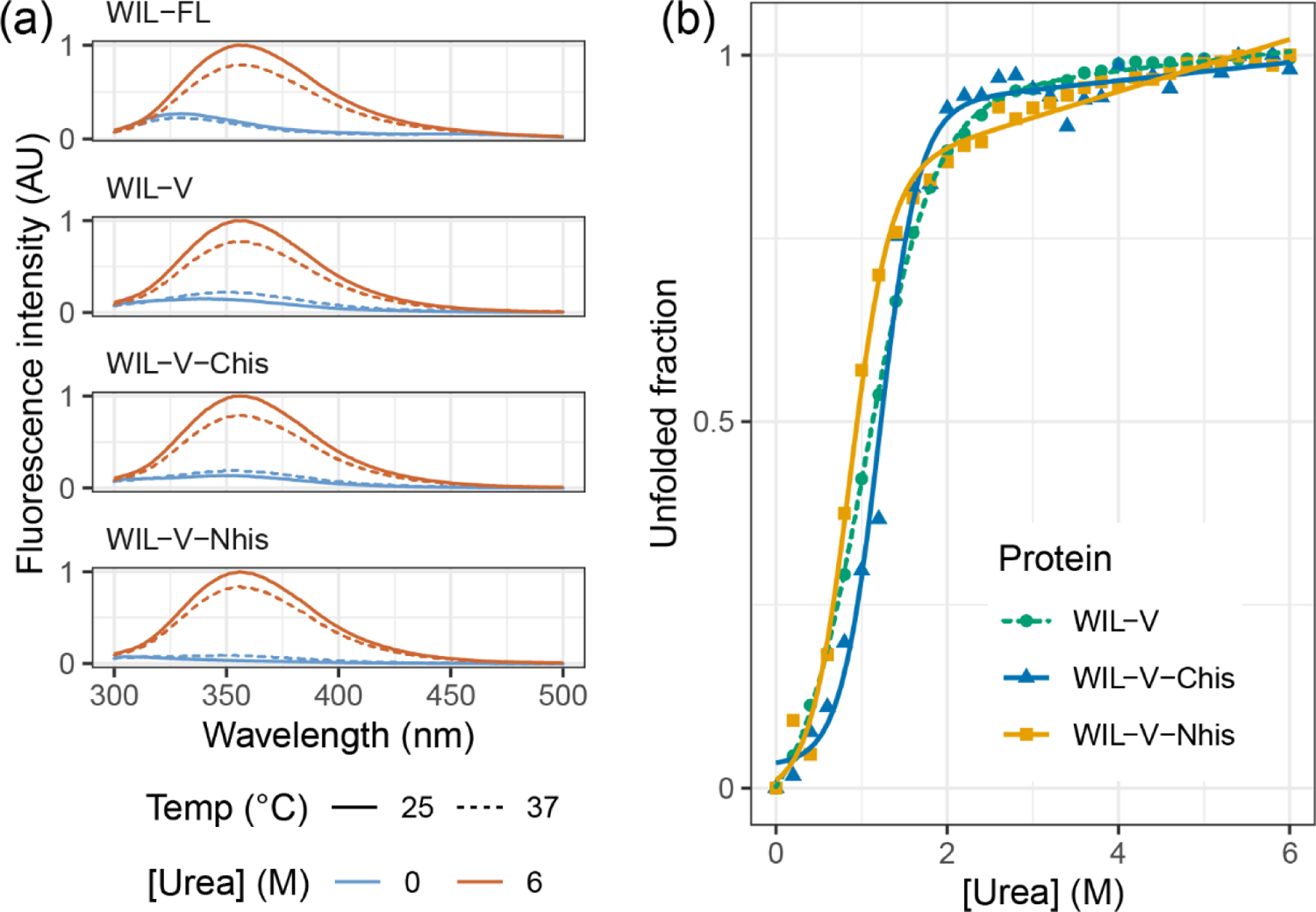
WIL-variants have similar folded states and stability. (a) Intrinsic fluorescence spectra (λex = 280 nm) of WIL variants showing increased fluorescence intensity and red-shifted fluorescence maxima upon unfolding in 6 M urea, which is characteristic of antibody variable domains. Data are shown at 25 °C and 37°C, consistent with the LCs remaining folded at 37 °C. (b) Urea titration of WIL-V variants in PBS, pH7.4, 37 °C, followed by intrinsic fluorescence. Colors and shapes indicate the variant: green circles, WIL-V; blue triangles, WIL-V-Chis; orange squares, WIL-V-Nhis.

At WIL-V concentrations of 4 µM or below, all three variants aggregated with comparable kinetics, although the endpoint fluorescence of WIL-V-Nhis was lower than that of other variants (Figure 3). This observation rules out the possibility that only AL55-like fibril structures are formed *in vitro*, since these are not compatible with an N-terminal extension. However, at higher concentrations the proteins’ T50 values diverged. The lag phase of WIL-V-Nhis increased by more than that of WIL-V. In contrast, the lag phase of WIL-V-Chis remained similar between 8 and 32 µM. These results are consistent with a model where off-pathway species are disrupted by the presence of a his-tag at the C-terminus but not the N-terminus of the V-domain.

**Figure 3:**
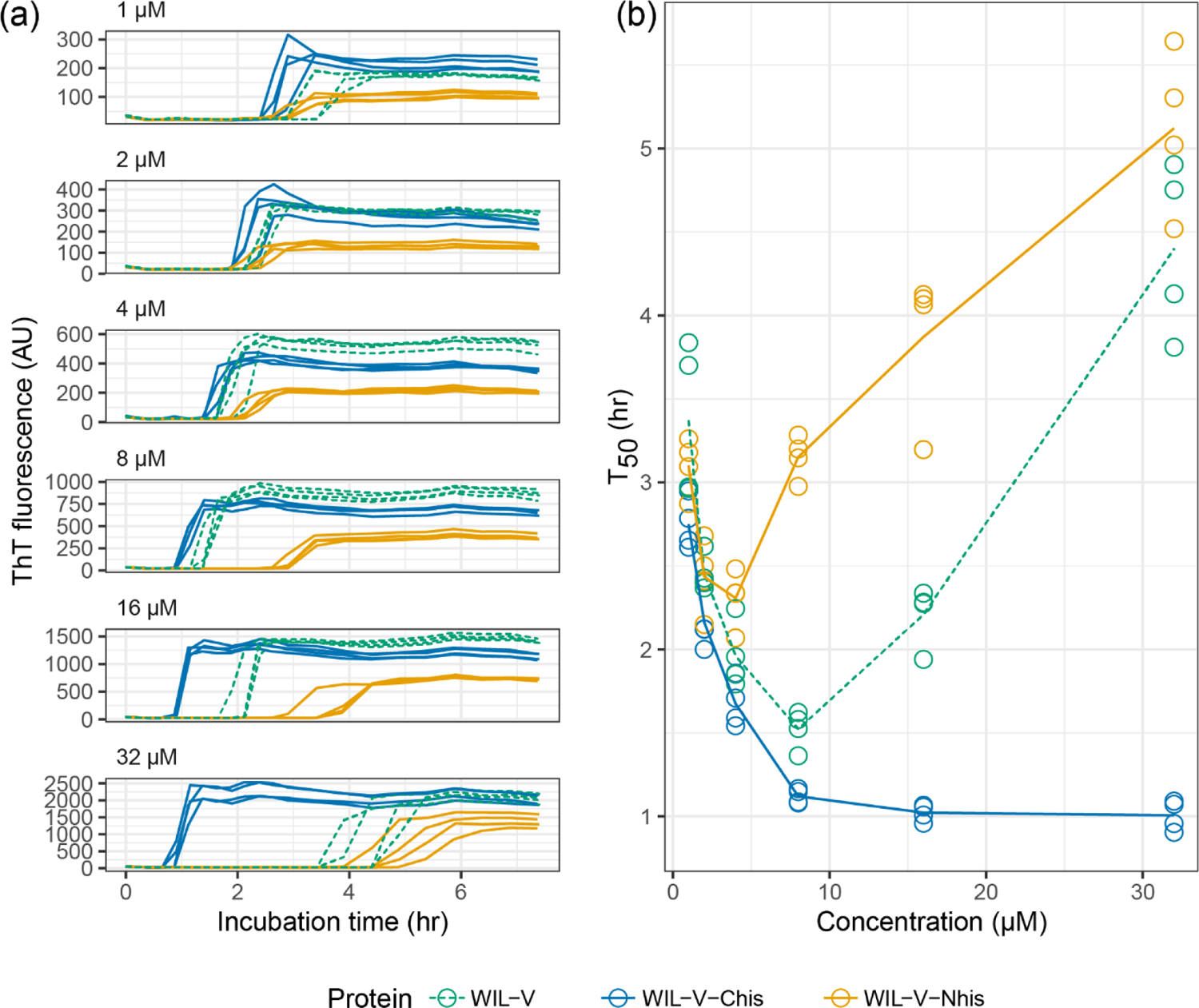
WIL-V variants aggregate with distinct kinetics. WIL-V solutions were incubated in microplates in PBS, pH 7.4, containing 1 µM ThT at 37 °C and shaken at 500 rpm. Data shown from two independent microwell plates, each with two wells per condition. The data for untagged WIL-V are reproduced from Figure 1. Colors are the same as in Figure 2. (a) ThT fluorescence as a function of time and concentration. (b) Calculated midpoint times (T_50_) for the data shown in (a). Lines show the average T_50_ at each timepoint.

One potential explanation for this behavior is a population of soluble, non-ThT-binding oligomers in solution, whose dissociation limits the rate of amyloid formation. To test whether such oligomeric species are present before the start of the reaction and could alter aggregation rates, we compared aggregation of WIL-V-Nhis before and after ultracentrifugation of the protein solution. Extended centrifugation clears particles larger than WIL-V dimers, which have sedimentation coefficients of around 2-3 Svedberg units. This treatment did not alter the aggregation kinetics of WIL-V-Nhis, so we conclude that large soluble species are not present at the start of aggregation reactions (Supplemental Figure 3).

### Fibril seeds have a limited ability to accelerate WIL-V aggregation

To investigate the ability of pre-formed aggregates to accelerate WIL-V aggregation, we added aggregated material (1% v/v) to new reactions and measured aggregation kinetics using ThT fluorescence. For all WIL-V variants, addition of seeds decreased the lag phase (Figure 4). However, the lag phases were not completely eliminated and the apparent rate at which ThT increased remained very similar between seeded and unseeded reactions. Titration of preformed Wil-V-Nhis seeds into WIL-V-Nhis aggregation reactions revealed that the lowest concentration used, 0.016% v/v or 1 in 6400, was sufficient to accelerate aggregate formation (Figure 5). However, addition of further seeds had only a limited effect. This saturation of seeding effect was observed at both 8 µM and 16 µM soluble protein concentration.

**Figure 4:**
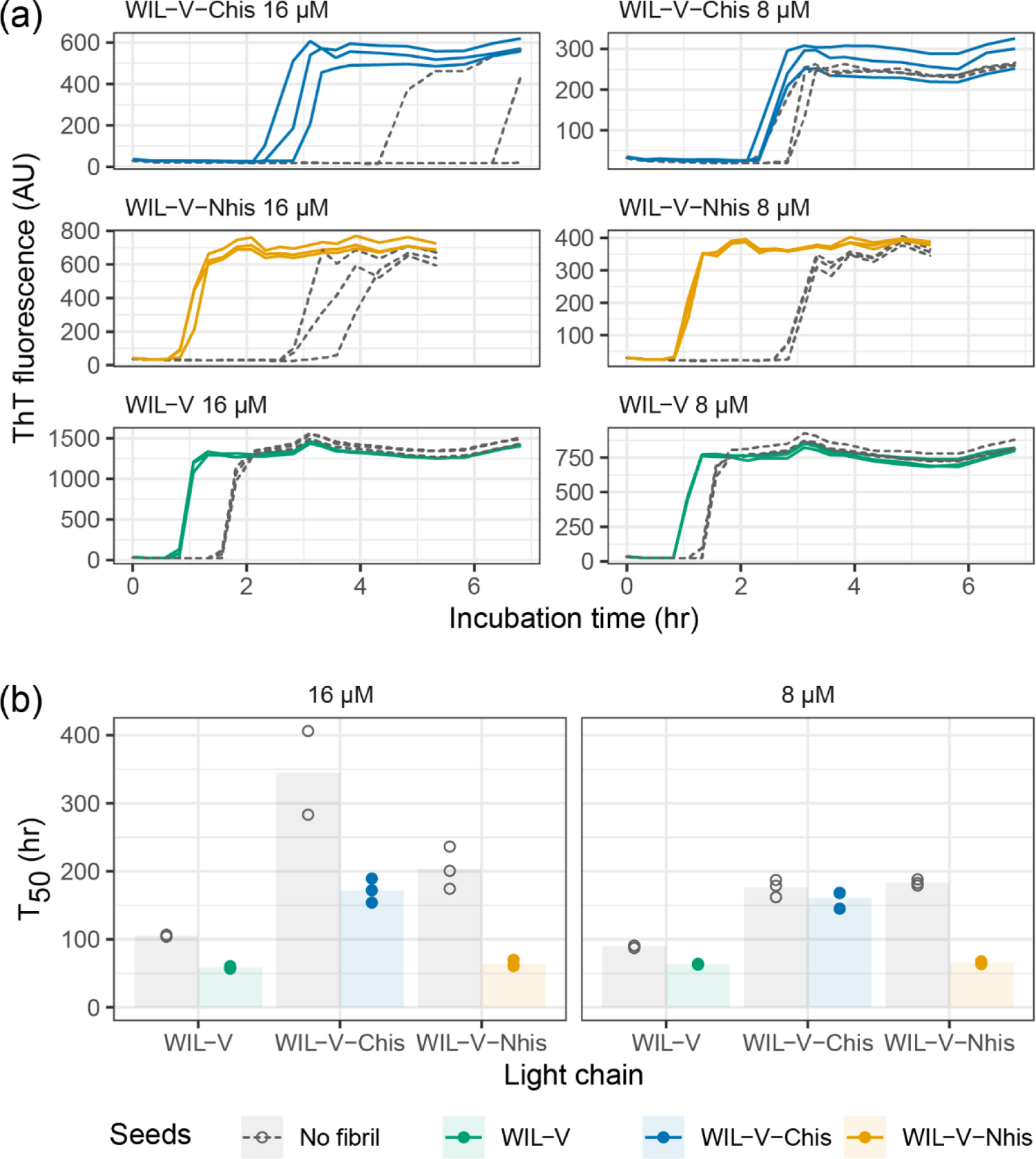
Aggregation of WIL-V variants can be accelerated by addition of homologous pre-formed seeds. WIL-V solutions were incubated in microplates, with (solid lines) or without (dashed lines) addition of 1% (v/v) pre-formed seeds (PBS, pH 7.4, 1 µM ThT, 37 °C, 500 rpm). Colors indicate the identity of the seed protein. (a) ThT fluorescence as a function of time. (b) Calculated midpoint times (T_50_) for the data shown in (a).

**Figure 5:**
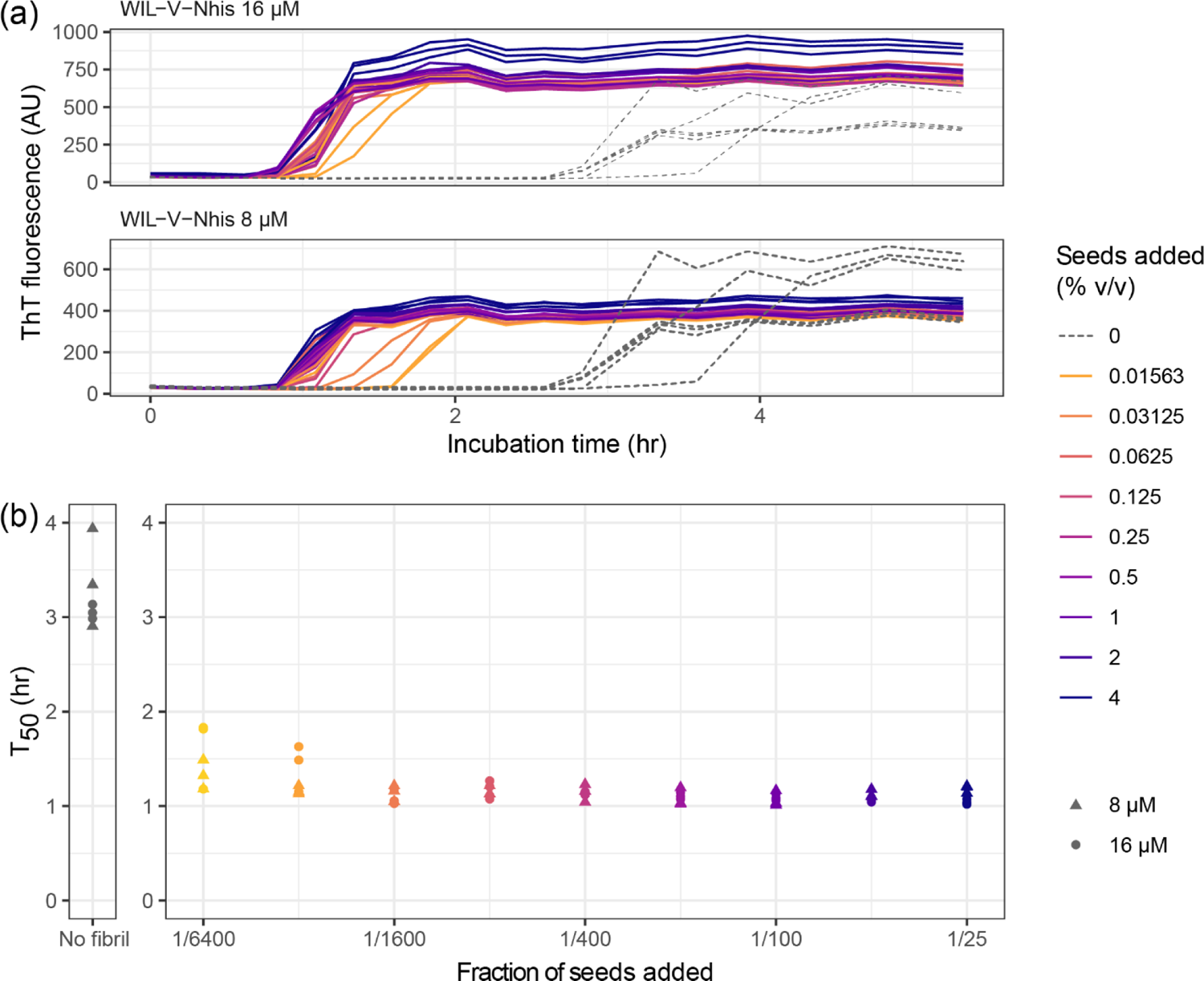
Concentration dependence of seeds on the aggregation kinetics of WIL-V-Nhis. Aggregation of WIL-V-Nhis in microplates (PBS, pH 7.4, 1 µM ThT, 37 °C, 500 rpm) was monitored by ThT fluorescence after addition of varying concentrations of pre-formed fibril seeds. Colors indicate the proportion of seeds added. Unseeded reactions are shown with dashed lines. (a) ThT fluorescence as a function of time and seed concentration. (b) Calculated midpoint times (T_50_) for the data shown in (a). Circles indicate T_50_ values observed at 8 µM, triangles indicate T_50_ values observed at 16 µM.

Finally, we asked whether pre-formed aggregates from one variant could accelerate aggregation of other variants. Addition of heterologous seeds reduced the lag phase of all variants at an initial soluble protein concentration of 32 µM, but at 8 µM only the effect on WIL-V-Nhis, which had the slowest aggregation, was consistent between replicates (Figure 6). No pre-formed aggregates of WIL-V variants were able to cause aggregation of WIL-FL within the 72 h experimental timecourse (Supplemental Figure 4).

**Figure 6:**
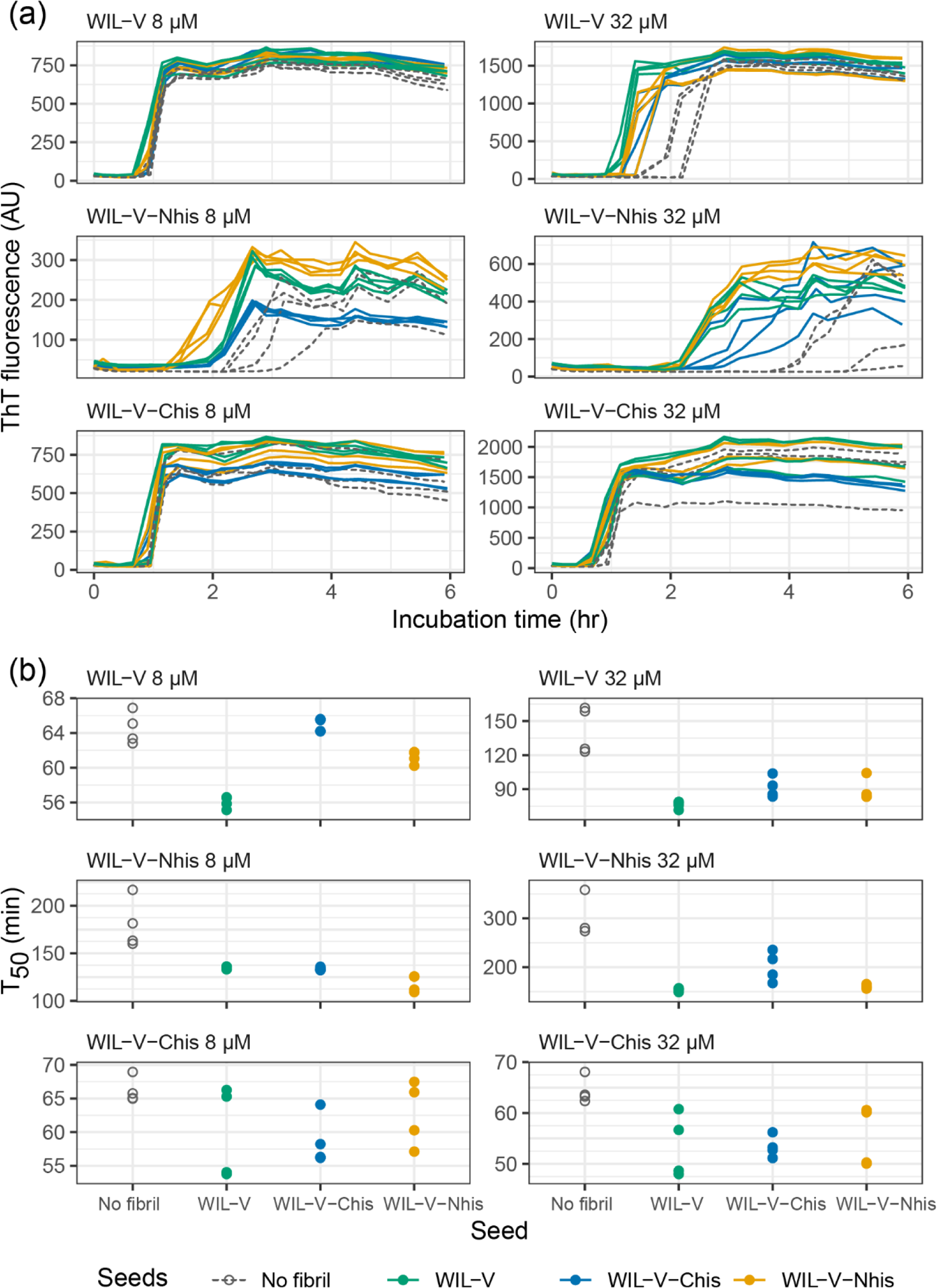
Heterologous seeds accelerate WIL-V aggregation with varying efficiency. Aggregation reactions in microplates were initiated in the presence or absence of 1% (v/v) seeds (PBS, pH 7.4, 1 µM ThT, 37 °C, 500 rpm). Colors indicate the identity of the seed protein. (a) ThT fluorescence as a function of time. Unseeded reactions are shown with dashed lines. (b) Calculated midpoint times (T_50_) for the data shown in (a).

## DISCUSSION

We investigated the aggregation mechanisms of the variable domain of the antibody light chain WIL, which was identified in a patient with AL amyloidosis. Under conditions where the protein is primarily folded (37 °C in PBS at pH 7.4, Figure 3a), morphologically diverse ThT-binding aggregates form rapidly after a lag phase of 1-4 hours (Figure 1). Increasing the concentration of WIL-V from 1 µM to 8 µm decreased the lag phase, but the lag phase increased at higher WIL-V concentrations. Variants of WIL-V with N- or C-terminal his-tags exhibited different aggregation kinetics that were not explained by differences in stability (Figure 4). Addition of pre-formed fibrils reduced, but did not eliminate the lag phase (Figure 5), and the tagged variants’ aggregation rates were similarly affected by the presence of homotypic or heterotypic seeds (Figure 6).

Previous observations, including our own experiments, observed that at a fixed concentration, the aggregation of light chain variable domains with similar sequences is highly correlated with their thermodynamic stability (Hurle et al., 1994; Wall et al., 1999; Souillac et al., 2002a; Rennella et al., 2019b). Consistent with these experiments, at low concentrations the presence of a his-tag does not greatly alter the aggregation rates observed for WIL-V variants. Instead, an N- or C-terminal his-tag appears to enhance or suppress, respectively, the formation of species that do not bind to ThT. Fibrils formed by WIL-V-Nhis cannot have the amyloid fold that was observed in the only IGLV6-57 fibrils reported to date (Swuec et al., 2019; Puri et al., 2023). Therefore, other cross-β fibril folds must be present. Furthermore, the observation of cross-seeding between WIL-V species means that mechanisms for acceleration of aggregation beyond exact templating of cross-β structures must be occurring. Such mechanisms may include “secondary” nucleation (Meisl et al., 2014), or propagation of a short, structured core of residues with the remainder of the protein sequence remaining unstructured, as has been proposed for.

These data highlight the complexity of aggregation mechanisms for initially-folded protein, compared to those for unstructured peptides such as Aß (Meisl et al., 2014, 2016). The extension of nuclei is generally rapid compared to initial formation of those nuclei for unstructured peptides. In consequence, both the lag phase and rate of aggregation during the growth phase have a clear dependence on protein concentration. Such systems typically have lag phases that can be eliminated by the addition of pre-formed fibrils. Aggregation of WIL-V under the conditions used here differs from these systems in at least three ways. First, we observed slow aggregation at increased protein concentration, which is consistent with the formation of off-pathway species that do not bind ThT. Second, ThT fluorescence increases rapidly during the growth phase. This is not consistent with the presence of off-pathway species, which might be expected to reduce the rate of fibril elongation as well as nucleation. We are not yet able to resolve this apparent contradiction. Third, pre-formed seeds have a limited capacity to accelerate aggregation and this seeding effect rapidly becomes saturated.

Thioflavin T fluorescence is widely used as a probe for cross-ß structure. Although we observe a rapid increase in ThT fluorescence and the signal intensities of the plateaus reached were proportional to the concentration of protein, the aggregates observed in these reactions were a mix of small fibrils and amorphous species. We previously observed much longer fibrils made by WIL-V and other LC proteins (Morgan and Kelly, 2016; Rennella et al., 2019b, 2019a). We were unable to observe differences in morphology between the WIL-V variants that might provide additional information about their structural differences. Similar results were observed in the aggregation of the FOR005 variable domain. Here, a rapid initial rise in ThT fluorescence was associated with the formation of small amorphous aggregates or “oligomers”, while fibrils appeared more slowly. It is possible that LCs rapidly form a subpopulation of small species that bind ThT more tightly than the “mature” fibrils. In contrast to the behavior of the WIL variable domain, we have previously observed heating full-length light chains to 55 °C led to formation of amorphous aggregates that did not bind to ThT (Cooley et al., 2014; Morgan and Kelly, 2016).

Early events in AL amyloid formation have been described for multiple light chains, revealing a diverse range of behavior (Souillac et al., 2002a, 2002b; Misra et al., 2019; Kazman et al., 2021; Misra and Ramirez-Alvarado, 2021; Pradhan et al., 2023). The rate at which variable domains aggregate *in vitro*, and the details of how loss of soluble protein correlates with an increase in ThT fluorescence vary substantially between light chains and between researchers. For example, two groups reported aggregation lag times for a lambda light chain variable domain known as FOR005 of four hours and four days at concentrations of 15 and 25 µM, respectively (Kazman et al., 2021; Pradhan et al., 2023). Several studies have identified non-fibrillar aggregate species, although the relationships between the various morphologies are not consistent between different reports. Furthermore, the kinetics of aggregation as measured by loss of soluble protein do not always correspond to an increase in ThT fluorescence.

Two studies by Ramirez-Alvarado and coworkers investigated the concentration dependence of aggregation and seeding behavior of AL-associated kappa variable domains and full-length light chain (Blancas-Mejía and Ramirez-Alvarado, 2016; Blancas-Mejía et al., 2017). Kappa light chains typically aggregate less readily than lambda light chains, which may be related to their under-representation in AL amyloidosis. These studies were carried out at pH 2, where the light chains readily formed amyloid fibrils on a timescale of hours to days. Similarly to the data observed for WIL-V, the kappa light chain variable domains had complex aggregation kinetics, in some cases with longer lag times at higher concentrations. Seeding, and in some cases heterologous seeding with other light chains, reduced the lag phase, but did not eliminate it, similar to the behavior observed for WIL-V. Pre-formed aggregates formed by variable domains were also able to accelerate aggregation of full-length light chains. Different behavior was observed in a study of variable domains with sequences corresponding to light chain germline precursor proteins (Garay Sánchez et al., 2017). Here, seeded aggregation occurred at 37 °C in PBS without agitation upon addition of 10% homologous seeds to 41.5 µM soluble LCs. In contrast to our results, ThT fluorescence and insoluble protein fraction increased in a near-linear manner over the course of several hours, without a lag phase.

An important limitation of this study is the method used to induce aggregation. Although the LC proteins are incubated under native-like conditions, reproducible aggregation kinetics can only be achieved by in our hands by vigorous shaking, outside the platereader. Agitation has multiple effects on protein aggregation, and the way that reactions are stirred or shaken can have substantial effects on the kinetics. Agitation can induce fragmentation of fibrils via shear forces to create additional ends, thereby increasing the elongation rate. Agitation can also alter the interactions between the reaction solution and its surroundings, the walls of the vessel (here, the polystyrene microwell plate) and the air. These surface effects are poorly understood. Identifying and accounting for these interactions, which may have analogs in patients but clearly cannot occur in the same way, will require further work.

Our investigations into the mechanisms of WIL-V aggregation have identified complex aggregation mechanisms that cannot readily be described by existing models. Using additional probes of amyloid structure may reveal more mechanistic details of how WIL-V and other LCs aggregate under native-like conditions. Aggregation *in vitro* can only capture a fraction of the complexity of amyloidosis in patients, where light chains also interact with a plethora of other molecules. However, differences in aggregation behavior between light chains may allow features that contribute to pathology to be identified. The data here highlight the importance of evaluating aggregation at multiple protein concentrations, both to optimize experimental conditions and determine how different sequence factors alter light chain aggregation.

## MATERIALS AND METHODS

### Materials

Chemicals were obtained from Thermo Fisher Scientific unless otherwise noted. Experiments were carried out in phosphate buffered saline at pH 7.4 (PBS): 10 mM Na_2_HPO_4_, 1.8 mM KH_2_PO_4_, 137 mM NaCl, and 2.7 mM KCl. Data analysis was carried out using the Tidyverse suite of tools in R via the RStudio environment (Wickham et al., 2019; R Core Team, 2020; RStudio Team, 2020), or in Wolfram Mathematica v.13 (Wolfram Research).

### Cloning and expression of WIL variants

WIL-V was previously cloned into a pET-22 expresion vector (EMD Millipore, USA) that encodes a C-terminal histidine tag after a TAA stop codon (Rennella et al., 2019a). Removal of this codon by site-directed mutagenesis (Q5 kit, New England Biolabs) yielded the WIL-V-Chis. To make WIL-V-Nhis, the sequence was cloned into a pQE2 expression vector (Qiagen) that encodes an N-terminal his-tag. WIL-V and WIL-V-Chis pET-22 constructs were transformed into RosettaGami (DE3) *E. coli* (EMD Millipore). The WIL-V-Nhis pQE2 construct was transformed into RosettaGami 2 *E. coli*. Cells transformed with WIL-V constructs were grown in lysogeny broth (LB) containing 100 µg/ml carbenicillin at 37 °C to an optical density (OD) of 0.6, cooled to 12 °C, induced with 125 µM isopropyl thiogalactopyranoside (IPTG) and expressed overnight. For WIL-FL, a pET-3 construct was transformed into BL21 (DE3) E. coli, grown in LB containing 100 µg/ml carbenicillin to an OD of 0.6 at 37 °C, induced with 125 µM IPTG and induced overnight at 37 °C in order to form inclusion bodies. Cells were harvested by centrifugation. DNase I (New England Biolabs) and 1 mM phenylmethyl sulfonyl fluoride were added and cells were lysed by osmotic shock (WIL-V and WIL-V-Chis) or sonication (WIL-V-Nhis and WIL-FL).

Purification of untagged WIL-V and WIL-FL was carried out as previously described (Morgan and Kelly, 2016). WIL-FL inclusion bodies were washed in PBS, resuspended in 4 M guanidine hydrochloride and dithiothreitol, then refolded dropwise into 25 mM Tris-Cl at pH 8.5, 4 °C. Refolded WIL-FL and untagged WIL-V were purified by ammonium sulfate precipitation, anion exchange and size-exclusion chromatography.

WIL-V-Chis and WIL-V-Nhis were loaded onto a Ni-NTA column (Cytiva HisTrap) in 2X PBS (20 mM Na_2_HPO_4_, 3.6 mM KH_2_PO_4_, 274 mM NaCl, and 5.4 mM KCl) containing 20 mM imidazole and eluted with a linear imidazole gradient (elution buffer was 2X PBS containing 500 mM imidazole). Eluted protein was concentrated and purified by size exclusion chromatography on a Superdex 75 column (Cytiva).

### Amyloid formation reactions

Aggregation experiments were run in 96 well clear bottom, black-walled plates (Corning #3631). Each experimental well contained 100 µL of filtered protein in 1x PBS with 1 µM ThT. Plates were sealed black vinyl film (Nunc, #236703) and covered with a lid to minimize evaporation. Protein serial dilutions from 32 µM to 1 µM concentration were used. Fluorescence was measured at 37 °C with a SpectraMax M5 platereader (Molecular Devices) using an excitation wavelength of 440 nm and emission wavelength of 480 nm. Plates were incubated at 37°C and shaken at approximately 500 rpm between readings (the plate shaker does not have a speed display).

Sedimentation of high molecular weight species before aggregation was carried out using a Type 50.2 Ti fixed-angle rotor in an Optima Ultracentrifuge (Beckman Coulter). Samples were centrifuged at 40,000 rpm (average 146,000 *g*), 4 °C for 30 h.

After unseeded fibril growth had reached an endpoint, samples from wells where the ThT fluorescence had increased were used to seed new reactions. The seeds from each reaction were added to new wells where the starting concentration of WIL-V was the same as that used for the initial unseeded reaction. Wells were mixed by pipetting and an appropriate volume was transferred to a new plate, so seed concentrations are expressed as volume ratios. For reactions where a fixed concentration of 1% seeds was used, 1 µl of seeds was added to a fresh reaction with a volume of 100 µl. For the seeding titration shown in Figure 5, seeds were first serially diluted into PBS before a fixed volume was transferred into the new wells. ThT fluorescence data were fit to a sigmoidal equation, using Wolfram Mathematica, to extract T_50_ values:

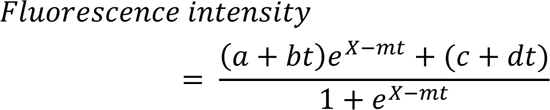

Where *a* and *c* are the intercepts of the post- and pre-transition baselines, respectively; *b* and *d* are the gradients of the post- and pre-transition baselines, respectively; *t* is the elapsed time; and *X* and *m* are constants describing the midpoint and slope of the transition. This equation is based on that describing a 2-state unfolding transition, which is not physically relevant but captures the shape of the ThT fluorescence curves to reliably calculate a midpoint time. The T_50_ values were calculated from the fit using the equation:

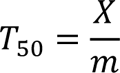

The AmyloFit website (https://amylofit.com/) was used as directed (Meisl et al., 2016). ThT fluorescence values were taken to correspond to relative aggregate concentration, and normalized using the website’s data entry tools. Triplicate data at concentrations of 1, 2, 4, 8, 16 and 32 µM were combined. Fits were attempted using the available models, but no satisfactory fit was identified.

### Fluorescence spectroscopy and urea titrations

Protein stability was evaluated using urea denaturation curves as previously described (Morgan and Kelly, 2016). Proteins at 5 µM concentration were titrated with 0 – 6 M urea in increments of 0.2 M urea in PBS at pH 7.4. Spectra was measured using a Fluoromax+ spectrometer (Horiba) with excitation at 280 nm and emission range of 300 – 500 nm at 25°C and 37°C. Spectra were compared using the intensity-weighted average wavelength of the emission spectrum, <λ>:

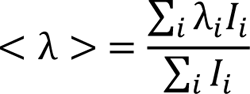

Where λi and Ii are the wavelength and intensity, respectively, at each measured point. The fraction of folded protein was calculated by normalizing the average wavelength to that of the 0 M and 6 M urea spectra. Titration data were fit to a two-state unfolding model for visual comparison, as previously described (Morgan and Kelly, 2016), but we did not attempt to quantitatively compare the titrations because of the short pre-transition baselines observed (Figure 2).

### Electron microscopy

Samples were diluted 10-fold in 10 mM Tris-Cl pH 7.4 and adsorbed onto Formvar/carbon-coated copper grids, stained with 1% uranyl acetate and washed with Tris-Cl. Grids were imaged using a CM12 transmission electron microscope (Philips Electron Optics) operated at 100 kV, and recorded with a Tietz 2Kx2K CCD camera (TVIPS) at 3 nm/pixel.

## ACKNOWLEDGEMENTS

We thank Dr. Elena Klimtchuk for help with protein expression and helpful discussions; Drs. Olga Gursky and Shobini Jayaraman for use of the ultracentrifuge; and Dr. Ester Bullitt for preparation of the electron micrographs. This work was supported by the Boston University Amyloidosis Research Fund and the Wildflower Foundation. MEW was supported by the Boston University Summer Training as Research Scholars Program, which is funded by NIH award R25HL118693. The authors declare no competing interests.

## SUPPLEMENTAL FIGURES

**Supplemental figure 1:**
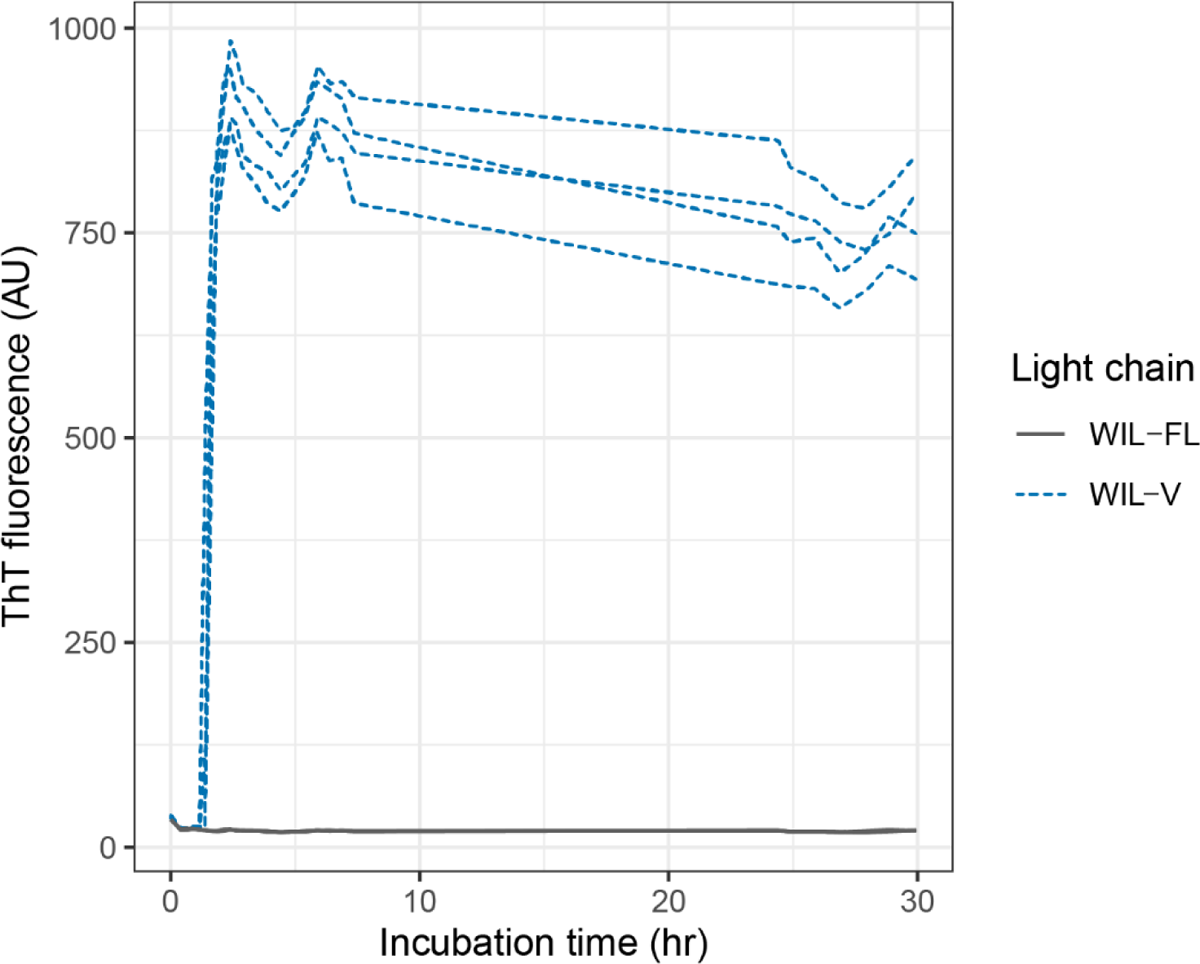
Full-length WIL remains soluble under conditions where its variable domain aggregates. WIL variants (n=4 wells) were incubated in microplates in PBS, pH 7.4, containing 1 µM ThT at 37 °C and shaken at 500 rpm. Aggregation kinetics were monitored by ThT fluorescence.

**Supplemental figure 2:**
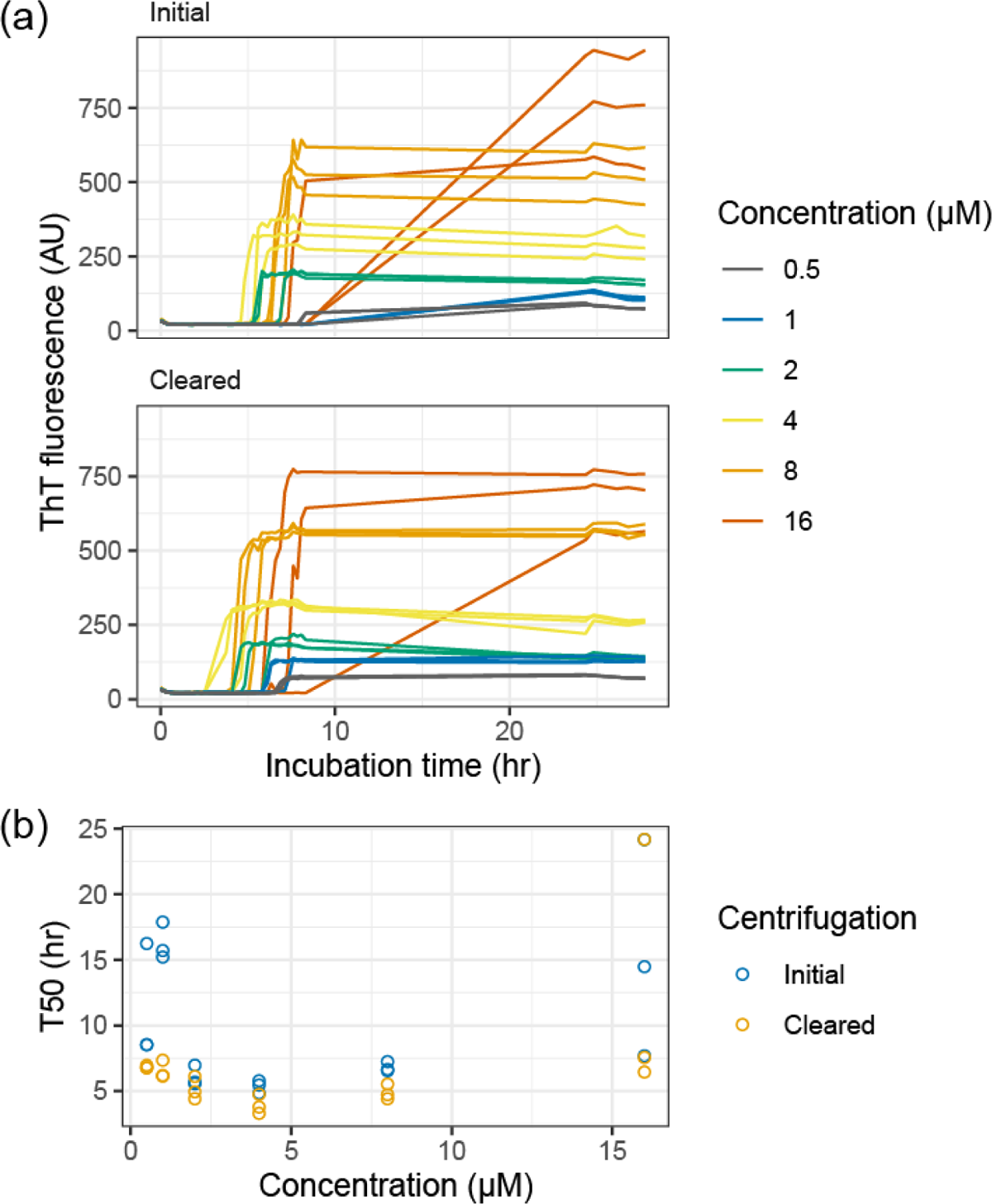
Pre-clearance of WIL-V-Nhis solutions by ultracentrifugation does not eliminate off-pathway aggregation. WIL-V-Nhis aggregation in PBS, with or without pre-clearance of the stock solution by ultracentrifugation, was monitored on a single microwell plate (n = 3 wells per condition) at 37 °C, 500 rpm. Note that slow aggregation occurred overnight when fluorescence could not be monitored for some samples, which limits the precision of these measurements. (a) ThT fluorescence as a function of time. Colors indicate initial protein concentrations. (b) Calculated midpoint times (T_50_) for the data shown in (a). Colors indicate pre-treatment.

**Supplemental figure 3:**
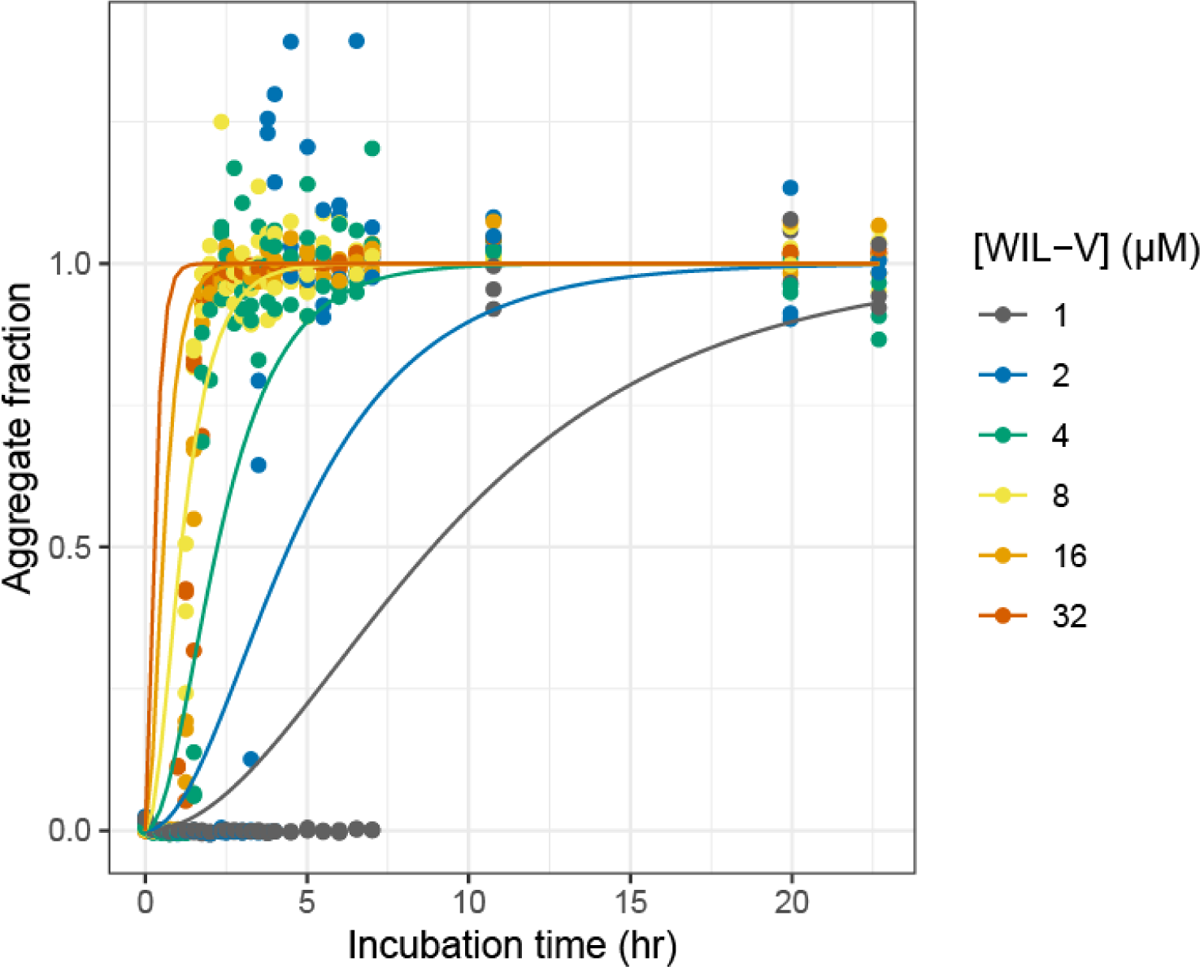
Existing models do not capture the kinetics of WIL-V aggregation. Example global fit of WIL-V aggregation kinetic data using the AmyloFit program’s nucleated polymerization model (Meisl et al., 2016). The model does not capture the long lag phase or rapid extension phase of aggregation, consistent with the hypothesis that more complex processes are involved.

**Supplemental Figure 4:**
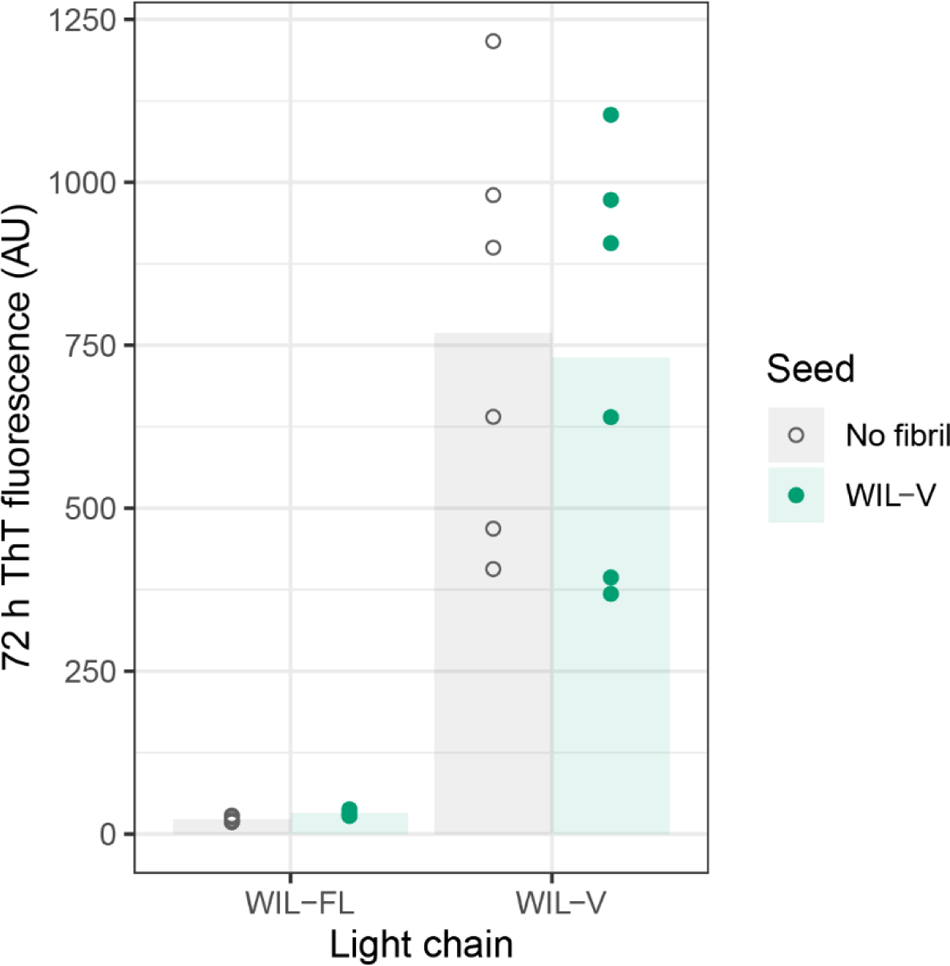
Addition of pre-formed fibril seeds does not lead to aggregation of full-length WIL within the timescale of these experiments. The ThT fluorescence observed in WIL-FL solutions (n = 6 wells) did not increase after 72 h incubation (8 µM LC, PBS, pH 7.4, 1 µM ThT, 37 °C, 500 rpm).

## Notes

### Competing Interest Statement

The authors have declared no competing interest.

## BIBLIOGRAPHY

Blancas-Mejía, L. M.; Ramirez-Alvarado, M. Recruitment of Light Chains by Homologous and Heterologous Fibrils Shows Distinctive Kinetic and Conformational Specificity. Biochemistry 2016, 55 (21), 2967–2978.

Blancas-Mejía, L. M.; Misra, P.; Ramirez-Alvarado, M. Differences in Protein Concentration Dependence for Nucleation and Elongation in Light Chain Amyloid Formation. Biochemistry 2017, 56 (5), 757–766.

Bodi, K.; Prokaeva, T.; Spencer, B.; Eberhard, M.; Connors, L. H.; Seldin, D. C. AL-Base: A Visual Platform Analysis Tool for the Study of Amyloidogenic Immunoglobulin Light Chain Sequences. Amyloid 2009, 16 (1), 1–8.

Buxbaum, J. N.; Dispenzieri, A.; Eisenberg, D. S.; Fändrich, M.; Merlini, G.; Saraiva, M. J. M.; Sekijima, Y.; Westermark, P. Amyloid Nomenclature 2022: Update, Novel Proteins, and Recommendations by the International Society of Amyloidosis (ISA) Nomenclature Committee. Amyloid 2022, 1–7.

Connors, L. H.; Jiang, Y.; Budnik, M.; Théberge, R.; Prokaeva, T.; Bodi, K. L.; Seldin, D. C.; Costello, C. E.; Skinner, M. Heterogeneity in Primary Structure, Post-Translational Modifications, and Germline Gene Usage of Nine Full-Length Amyloidogenic Kappa1 Immunoglobulin Light Chains. Biochemistry 2007, 46 (49), 14259–14271.

Cooley, C. B.; Ryno, L. M.; Plate, L.; Morgan, G. J.; Hulleman, J. D.; Kelly, J. W.; Wiseman, R. L. Unfolded Protein Response Activation Reduces Secretion and Extracellular Aggregation of Amyloidogenic Immunoglobulin Light Chain. Proc. Natl. Acad. Sci. U. S. A. 2014, 111 (36), 13046–13051.

Enqvist, S.; Sletten, K.; Stevens, F. J.; Hellman, U.; Westermark, P. Germ Line Origin and Somatic Mutations Determine the Target Tissues in Systemic AL-Amyloidosis. PLoS One 2007, 2 (10), e981.

Enqvist, S.; Sletten, K.; Westermark, P. Fibril Protein Fragmentation Pattern in Systemic AL-Amyloidosis. J. Pathol. 2009, 219 (4), 473–480.

Feige, M. J.; Hendershot, L. M.; Buchner, J. How Antibodies Fold. Trends Biochem. Sci. 2010, 35 (4), 189–198.

Ferrone, F. Analysis of Protein Aggregation Kinetics. Methods Enzymol. 1999, 309, 256–274.

Garay Sánchez, S. A.; Rodríguez Álvarez, F. J.; Zavala-Padilla, G.; Mejia-Cristobal, L. M.; Cruz-Rangel, A.; Costas, M.; Fernández Velasco, D. A.; Melendez-Zajgla, J.; Del Pozo-Yauner, L. Stability and Aggregation Propensity Do Not Fully Account for the Association of Various Germline Variable Domain Gene Segments with Light Chain Amyloidosis. Biol. Chem. 2017, 398 (4), 477–489.

Gorski, S. A.; Capaldi, A. P.; Kleanthous, C.; Radford, S. E. Acidic Conditions Stabilise Intermediates Populated during the Folding of Im7 and Im9. J. Mol. Biol. 2001, 312 (4), 849–863.

Hurle, M. R.; Helms, L. R.; Li, L.; Chan, W.; Wetzel, R. A Role for Destabilizing Amino Acid Replacements in Light-Chain Amyloidosis. Proc. Natl. Acad. Sci. U. S. A. 1994, 91 (12), 5446–5450.

Kazman, P.; Absmeier, R. M.; Engelhardt, H.; Buchner, J. Dissection of the Amyloid Formation Pathway in AL Amyloidosis. Nat. Commun. 2021, 12 (1), 6516.

Kourelis, T. V.; Dasari, S.; Theis, J. D.; Ramirez-Alvarado, M.; Kurtin, P. J.; Gertz, M. A.; Zeldenrust, S. R.; Zenka, R. M.; Dogan, A.; Dispenzieri, A. Clarifying Immunoglobulin Gene Usage in Systemic and Localized Immunoglobulin Light-Chain Amyloidosis by Mass Spectrometry. Blood 2017, 129 (3), 299–306.

Lavatelli, F.; Perlman, D. H.; Spencer, B.; Prokaeva, T.; McComb, M. E.; Théberge, R.; Connors, L. H.; Bellotti, V.; Seldin, D. C.; Merlini, G.;, et al. Amyloidogenic and Associated Proteins in Systemic Amyloidosis Proteome of Adipose Tissue. Mol. Cell. Proteomics 2008, 7 (8), 1570–1583.

Meisl, G.; Yang, X.; Hellstrand, E.; Frohm, B.; Kirkegaard, J. B.; Cohen, S. I. A.; Dobson, C. M.; Linse, S.; Knowles, T. P. J. Differences in Nucleation Behavior Underlie the Contrasting Aggregation Kinetics of the Aβ40 and Aβ42 Peptides. Proceedings of the National Academy of Sciences 2014, 111 (26), 9384–9389.

Meisl, G.; Kirkegaard, J. B.; Arosio, P.; Michaels, T. C. T.; Vendruscolo, M.; Dobson, C. M.; Linse, S.; Knowles, T. P. J. Molecular Mechanisms of Protein Aggregation from Global Fitting of Kinetic Models. Nat. Protoc. 2016, 11 (2), 252–272.

Merlini, G.; Dispenzieri, A.; Sanchorawala, V.; Schönland, S. O.; Palladini, G.; Hawkins, P. N.; Gertz, M. A. Systemic Immunoglobulin Light Chain Amyloidosis. Nat Rev Dis Primers 2018, 4 (1), 38.

Misra, P.; Ramirez-Alvarado, M. Early Events in Light Chain Aggregation at Physiological PH Reveal New Insights on Assembly, Stability, and Aggregate Dissociation. Amyloid 2021, 28 (2), 113–124.

Misra, P.; Blancas-Mejia, L. M.; Ramirez-Alvarado, M. Mechanistic Insights into the Early Events in the Aggregation of Immunoglobulin Light Chains. Biochemistry 2019, 58 (29), 3155–3168.

Morgan, G. J. Transient Disorder along Pathways to Amyloid. Biophys. Chem. 2022, 281, 106711.

Morgan, G. J.; Kelly, J. W. The Kinetic Stability of a Full-Length Antibody Light Chain Dimer Determines Whether Endoproteolysis Can Release Amyloidogenic Variable Domains. J. Mol. Biol. 2016, 428 (21), 4280–4297.

Morgan, G. J.; Buxbaum, J. N.; Kelly, J. W. Light Chain Stabilization: A Therapeutic Approach to Ameliorate AL Amyloidosis. Hematology 2021, 2 (4), 645–659.

Muchtar, E.; Dispenzieri, A.; Magen, H.; Grogan, M.; Mauermann, M.; McPhail, E. D.; Kurtin, P. J.; Leung, N.; Buadi, F. K.; Dingli, D.;, et al. Systemic Amyloidosis from A (AA) to T (ATTR): A Review. J. Intern. Med. 2021, 289 (3), 268–292.

Olsen, K. E.; Sletten, K.; Westermark, P. Fragments of the Constant Region of Immunoglobulin Light Chains Are Constituents of AL-Amyloid Proteins. Biochem. Biophys. Res. Commun. 1998, 251 (2), 642–647.

Powers, E. T.; Powers, D. L. Mechanisms of Protein Fibril Formation: Nucleated Polymerization with Competing off-Pathway Aggregation. Biophys. J. 2008, 94 (2), 379–391.

Pradhan, T.; Sarkar, R.; Meighen-Berger, K. M.; Feige, M. J.; Zacharias, M.; Reif, B. Mechanistic Insights into the Aggregation Pathway of the Patient-Derived Immunoglobulin Light Chain Variable Domain Protein FOR005. Nat. Commun. 2023, 14 (1), 3755.

Puri, S.; Schulte, T.; Chaves-Sanjuan, A.; Mazzini, G.; Caminito, S.; Pappone, C.; Anastasia, L.; Milani, P.; Merlini, G.; Bolognesi, M.;, et al. The Cryo-EM Structure of Renal Amyloid Fibril Suggests Structurally Homogeneous Multiorgan Aggregation in AL Amyloidosis. J. Mol. Biol. 2023, 168215.

R Core Team. R: A Language and Environment for Statistical Computing https://www.R-project.org/.

Radamaker, L.; Lin, Y.-H.; Annamalai, K.; Huhn, S.; Hegenbart, U.; Schönland, S. O.; Fritz, G.; Schmidt, M.; Fändrich, M. Cryo-EM Structure of a Light Chain-Derived Amyloid Fibril from a Patient with Systemic AL Amyloidosis. Nat. Commun. 2019, 10 (1), 1103.

Radamaker, L.; Baur, J.; Huhn, S.; Haupt, C.; Hegenbart, U.; Schönland, S.; Bansal, A.; Schmidt, M.; Fändrich, M. Cryo-EM Reveals Structural Breaks in a Patient-Derived Amyloid Fibril from Systemic AL Amyloidosis. Nat. Commun. 2021a, 12 (1), 875.

Radamaker, L.; Karimi-Farsijani, S.; Andreotti, G.; Baur, J.; Neumann, M.; Schreiner, S.; Berghaus, N.; Motika, R.; Haupt, C.; Walther, P.;, et al. Role of Mutations and Post-Translational Modifications in Systemic AL Amyloidosis Studied by Cryo-EM. Nat. Commun. 2021b, 12 (1), 6434.

Rennella, E.; Morgan, G. J.; Kelly, J. W.; Kay, L. E. Role of Domain Interactions in the Aggregation of Full-Length Immunoglobulin Light Chains. Proc. Natl. Acad. Sci. U. S. A. 2019a, 116 (3), 854–863.

Rennella, E.; Morgan, G. J.; Yan, N.; Kelly, J. W.; Kay, L. E. The Role of Protein Thermodynamics and Primary Structure in Fibrillogenesis of Variable Domains from Immunoglobulin Light Chains. J. Am. Chem. Soc. 2019b, 141 (34), 13562–13571.

RStudio Team. RStudio: Integrated Development Environment for R. RStudio, PBC.: Boston, MA 2020.

Souillac, P. O.; Uversky, V. N.; Millett, I. S.; Khurana, R.; Doniach, S.; Fink, A. L. Effect of Association State and Conformational Stability on the Kinetics of Immunoglobulin Light Chain Amyloid Fibril Formation at Physiological PH. J. Biol. Chem. 2002a, 277 (15), 12657–12665.

Souillac, P. O.; Uversky, V. N.; Millett, I. S.; Khurana, R.; Doniach, S.; Fink, A. L. Elucidation of the Molecular Mechanism during the Early Events in Immunoglobulin Light Chain Amyloid Fibrillation. Evidence for an off-Pathway Oligomer at Acidic PH. J. Biol. Chem. 2002b, 277 (15), 12666–12679.

Swuec, P.; Lavatelli, F.; Tasaki, M.; Paissoni, C.; Rognoni, P.; Maritan, M.; Brambilla, F.; Milani, P.; Mauri, P.; Camilloni, C.;, et al. Cryo-EM Structure of Cardiac Amyloid Fibrils from an Immunoglobulin Light Chain AL Amyloidosis Patient. Nat. Commun. 2019, 10 (1), 1269.

Wall, J.; Schell, M.; Murphy, C.; Hrncic, R.; Stevens, F. J.; Solomon, A. Thermodynamic Instability of Human Lambda 6 Light Chains: Correlation with Fibrillogenicity. Biochemistry 1999, 38 (42), 14101–14108.

Wickham, H.; Averick, M.; Bryan, J.; Chang, W.; McGowan, L. D.; François, R.; Grolemund, G.; Hayes, A.; Henry, L.; Hester, J.;, et al. Welcome to the tidyverse. Journal of Open Source Software 2019, 4 (43), 1686.

